# KDM6A is enzymatically active at the Golgi apparatus where it is necessary for maintenance of Golgi apparatus morphology

**DOI:** 10.64898/2026.11.02.763691

**Authors:** Charli S. Worth, Jessica M. Conforti, Adhwaitha M. Nambiar, Jack Malone, Jacob T. Bailey, Rabin Buthodokhi, Amanda M. Ziegler, Kshetra Polavarapu, Hannah Lilley, Leslie K. Climer, Dwayne D Simmons, Jonathan A. Kelber, Elyssia S. Gallagher, Joseph H. Taube

**Affiliations:** Department of Biology, Baylor University, Waco, TX, USA; Department of Chemistry and Biochemistry, Baylor University, Waco, TX, USA

**Keywords:** Golgi, Lysine demethylases, EMT, KDM6A

## Abstract

KDM6A and KDM6B, lysine demethylases, play a central role in regulation of transcription through their demethylation activity towards H3K27me3 and have also been shown to be dynamically regulated upon epithelial-mesenchymal transition (EMT). In particular, KDM6A has been implicated in a variety of malignancies and is frequently mutated in cancers including those in kidney, bladder, colorectal, and breast. KDM6A and KDM6B function as part of the COMPASS-like protein complex (CLC) which includes methyltransferases KMT2C/KMT2D and the regulatory WRAD-subcomplex. KDM6A total protein is known to be suppressed by TGF-β in MCF10A breast cells; however, our examination of protein localization by immunofluorescence and LC-MS/MS reveals that KDM6A expression is lost in the nucleus but retained in the Golgi apparatus, independent of other CLC complex members. Importantly, KDM6A isolated from a biochemically-separated subcellular membranous fraction possesses the catalytic JmjC domain and is enzymatically active, indicating that KDM6A may have non-nuclear substrates. Further, knockdown of KDM6A and treatment with TGF-β similarly alter Golgi apparatus morphology, in a cell type specific manner, suggesting context-dependent regulatory functions. These findings support a model in which KDM6A is necessary to maintain Golgi apparatus morphology and is enzymatically active outside of the nucleus.

## INTRODUCTION

Post-translational modifications (PTMs) are covalent modifications to proteins that impact their structure, function, and interactions [1]. PTMs on histone proteins function as crucial epigenetic markers by altering gene expression through modifying chromatin structure or through recruitment of ‘reader’ proteins [2]. Methylation of lysine residues on histone tails is regulated by lysine methyltransferase (KMTs) and lysine demethylase proteins (KDMs) [3] [4]. Lysine 27 on histone H3 (H3K27) can be methylated up to three times (H3K27me3) by the SET-domain methyltransferase EZH2, the catalytic component of the PRC2 complex, and is associated with developmentally regulated gene silencing [5] [6]. Catalytic removal of H3K27 tri- or di-methylation by paralogous proteins KDM6A or KDM6B is accomplished by the catalytic Jumonji C domain (JmjC), requires O_2_, Fe (II), and alpha ketoglutarate (a-KG) co-factors, and terminates in a mono-methylated residue (H3K27me1) [7] [8].

KDM6A and KDM6B are known to function as part of the COMPASS-like protein complex (CLC), a variant of the COMPASS complex [9]. The CLC is a highly conserved, multi-protein complex. In addition to either KDM6A or KDM6B, its members include either of two H3K4 methyltransferase proteins KMT2C or KMT2D, plus the four-member WRAD subcomplex: WDR5, RBBP5, ASH2L, and DPY30. While both the CLC and the COMPASS complex include WRAD proteins, the COMPASS complex substitutes H3K4 methyltransferase core proteins KMT2C or KMT2D with either KMT2A or KMT2B [10] [7] [11]. Both complexes are involved in transcription activation through methylation of histone H3 lysine 4 (H3K4) and through the recruitment of lysine acetyltransferases [12, 13]. Though the functions of KDM6A and other proteins in the CLC are well characterized in their canonical roles on the chromatin fiber, potential non-nuclear functions of KDM6A are not well understood.

The Golgi apparatus plays a central role in the trafficking and sorting of proteins received from the endoplasmic reticulum. Additionally the Golgi apparatus facilitates addition of PTMs, particularly carbohydrate, phosphate, and sulfate groups [14]. Both expression of KDM6A and morphological changes to this organelle have been separately associated with epithelial-to-mesenchymal transition (EMT), a process which imparts the motility enabling migration during development or metastasis of cancer cells [15] [16]. However, to the best of our knowledge, neither function of KDM6 proteins outside the nucleus nor a connection between KDM6 proteins and Golgi apparatus morphology has been reported.

In this work, we detail KDM6A and KDM6B localization using multiple methodologies. We observe persistent localization of enzymatically active KDM6A to the Golgi apparatus despite EMT-associated suppression of KDM6A in the nucleus; and demonstrate that Golgi apparatus morphology is impacted by KDM6A. These data highlight a novel, non-nuclear role for lysine demethylases and point towards an expanded view of epigenetic regulation to include actions of lysine demethylases outside the nucleus.

## METHODS

### Cell culture

All cell lines, MCF7, MDA-MB-231, 293T, and MCF10A were purchased from ATCC (Manassas, VA, USA). HMLE, HMLE-Ras, and HMLE TWIST were kindly gifted from Dr. Sendurai Mani (MD Anderson Cancer, Center) and were cultured as previously described [17]. MCF7, MDA-MB-231, and 293T were cultured in Dulbecco’s Modified Eagle’s Medium (DMEM) (Corning Inc., Kennebuck, ME, USA) supplemented with 10% fetal bovine serum (FBS) (Equitech-Bio Inc., Kerrville, Texas, USA) and 1% antibiotics (Penicillin/Streptomycin, Lonza, Basel, Switzerland). MCF10A cells were cultured in DMEM:F12 media supplemented with 5% horse serum (HS) (Cytiva, Marolborough, MA, USA), 1% penicillin/streptomycin (Gibco, Billings, MT, USA), 500 ng/mL hydrocortisone (ACROS, Geel, Belgium), 100 ng/mL cholera toxin (Sigma-Aldrich, St. Louis, MO, USA), 0.01 mg/mL insulin (Sigma-Aldrich, St. Louis, MO, USA), and 20 ng/mL EGF (EMD Millipore Corp, Bulington, MA, USA). Cells were incubated at 37°C with 5% CO_2_ and passaged at least every third day to maintain consistent densities. Cells were regularly tested for mycoplasma contamination using PlasmoTest Mycoplasma Detection Kit (InvivoGen, San Diego, CA, USA).

### Chemical treatment

For TGF-β treatment, media was supplemented with 5.0 ng/mL recombinant human TGF-β-1 (R&D Systems, Minneapolis, MN, USA; resuspended in 4 mM HCl, 0.1% BSA) for the indicated amounts of time. Reconstitution solutions served as vehicle controls.

### Immunoblotting

Cells were washed with ice-cold PBS and RIPA buffer (Alfa Aesar, Haverhill, MA, USA), supplemented with protease and phosphatase inhibitors (Thermo Fisher Scientific, Waltham, MA, USA). Cells were collected using a cell scraper and incubated at 4 °C for 30 minutes. Lysed cells were then sonicated with a hand sonicator three times for two seconds each at 40% power with one minute reset in between before an additional incubation for 30 minutes at 4 °C. Lysates were centrifuged at 15,000 RCF for 20 min at 4 °C and the supernatant was recovered. Protein concentration was determined using a bicinchoninic acid assay (BCA) (Thermo Fisher Scientific, Waltham, MA, USA). Proteins were separated on a 10% SDS-PAGE gel and transferred onto a 0.45 µm PVDF membrane (Thermo Fisher Scientific, Waltham, MA, USA). After transfer, blots were blocked with 5% non-fat skim milk and probed with the appropriate primary and secondary antibodies. Chemiluminescence signal was obtained using ECLTM Prime (Cytiva, Marlborough, MA, USA), and blot images were captured using the ChemiDocTM Imaging System (Bio-Rad, Hercules, CA, USA). When necessary, blots were stripped with ECL Stripping Buffer (Li-Cor, Lincoln, NE, USA) following manufacturer protocol for reprobing.

### Immunofluorescence

Cells were seeded at 25,000 cells per coverslip in a 6 well plate two days prior to fixation. Cells were fixed in 2% paraformaldehyde (P1010, Specturm Chemical MGF Corp, Gardena, CA, USA) in PBS for 20 minutes and washed twice with PBS before permeabilization with 0.3% Triton X-100 (BP151-500, Thermo Fisher Scientific, Waltham, MA) in PBS for 15 minutes. Cells were washed twice with PBS before being quenched with 1% glycine (BP381, Themo Fisher Scientific, Waltham, MA) in PBS for 5 minutes followed by two PBS washes before being blocked with 8% BSA (BP1605-100, Fisher Bioreagents, Waltham, MA, USA) in TBST for overnight at 4 °C. Coverslips were incubated overnight with primary antibodies diluted in TBST at 4 °C in a humidity chamber followed by four TBST washes. This was followed by a one-hour incubation with secondary antibodies diluted in 3% BSA in TBST followed by four TBST washes.

Coverslips were then incubated for 5 minutes with DAPI (D1306, Thermo Fisher Scientific, Waltham, MA) before being washed once with water and mounted with permount (SP15-100, Thermo Fisher Scientific, Waltham, MA) to a microscope slide. Slides were stored at 4°C and imaged with an Agilent BioTek Cytation C10 confocal microscope.

Images were also acquired using a Zeiss LSM 800 upright laser scanning confocal microscope (Carl Zeiss Microscopy, Jena, Germany). Near super-resolution imaging was performed using the Airyscan detection module with a Plan-Apochromat 63×/1.4 NA oil-immersion DIC M27 objective. Z-stack image series were collected using optimized acquisition settings in the x, y, and z dimensions. Airyscan raw data were processed using the Airyscan reconstruction algorithm in Zeiss ZEN software (version 2.x, Carl Zeiss Microscopy) to generate super-resolution images with enhanced signal-to-noise ratio and spatial resolution. Identical laser power, detector gain, and processing parameters were maintained for all samples within each experimental comparison.

### Antibodies

The following antibodies were used: KDM6A (NBP1-80628, Novus Biologicals, Centennial, CO, USA), KDM6B (NBP1-06640, Novus Biologicals, Centennial, CO, USA), KMT2C (MA-38554, Invitrogen, Carlsbad, CA, USA), KMT2D (PA5-49581, Invitrogen, Carlsbad, CA, USA), WDR5 (703745, Invitrogen, Carlsbad, CA, USA), ASH2L (D93F6, Cell Signaling Technologies, Danvers, MA, USA), SP1 (D4C3, Cell Signaling Technologies, Danvers, MA, USA), GM130 (610823, BD Biosciences, Franklin Lakes, NJ, USA), Actin (8457S, Cell Signaling Technologies, Danvers, MA, USA), Actin (612657, BD Biosciences, Franklin Lakes, NJ, USA), GAPDH (21185, Cell Signaling Technologies, Danvers, MA, USA), H3 (9715S, Cell Signaling Technologies, Danvers, MA, USA), Vimentin (5741, Cell Signaling Technologies, Danvers, MA, USA), anti-rabbit IgG Fab2 Alexa Fluor ®488 (4412, Cell Signaling Technologies, Danvers, MA, USA), anti-rabbit IgG Fab2 Alexa Fluor ®594 (8889, Cell Signaling Technologies, Danvers, MA, USA), anti-mouse IgG Fab2 Alexa Fluor ®488 (4408, Cell Signaling Technologies, Danvers, MA, USA), and anti-mouse IgG Fab2 Alexa Fluor ®594 (8890, Cell Signaling Technologies, Danvers, MA, USA)

### Immunoprecipitation

MCF7 cells were fractioned via subcellular protein fractionation (78840, Thermo Fisher Scientific, Waltham, MA, USA) into cytoplasmic, membranous, soluble nuclear, and chromatin-bound fractions. Once lysates were obtained, 10 μL of KDM6A primary antibody (NBPI-80628, Novus Biologicals, Centennial, CO, USA) or 10 μL of Normal Mouse IgG (sc-2025, Santa Cruz, Dallas, TX, USA) was added to 1 mL of lysate and incubated at 4°C overnight. Protein A/G magnetic agarose beads (78609, Thermo Fisher Scientific, Waltham, MA, USA), were pre-washed and added to the lysate plus antibody solution post overnight incubation for 20 minutes with gentle rotation. Once the solution had finished mixing, beads were pelleted on a magnetic separation rack on ice and the supernatant was removed. The beads were washed five times with 250 μL of buffer (respective to the sample from subcellular protein fractionation kit buffer) and the supernatant was removed. The beads were washed with 500uL of wash buffer (0.1% SDS: 1% TRITON; 2 mM EDTA; 20 mM TRIS HCl pH 8) twice and supernatant was discarded. To recover bound proteins, 100 μL of elution buffer (10 mM TRIS pH 8.5 in H_2_O) was added to the beads and incubated for 10 minutes at room temperature with occasional mixing. The beads were collected with a magnetic stand and the supernatant was saved and stored at –80 °C until use.

### Generation of Knockdown Cell Lines

Plasmids (pGIPZ) for shRNA mediated suppression of KDM6A, and a scrambled control, were obtained from Horizon Discovery. Lentiviral transduction was conducted in 293T cells (ATCC) using FuGENE HD transfection reagent (E2311, Promega, Madison, WI). Viral supernatant was collected and filtered through a 0.45 μm filter (9915-2504, Cytiva, Marlborough, MA). Transduction to target cell lines was aided with 8 μg/mL protamine sulfate (9009-65-8, Thermo Fisher Scientific, Waltham, MA) and viral supernatant was allowed to incubate for 24 hours. Knockdown cell lines were selected via puromycin (ant-pr, Invivogen, San Diego, CA). Cell lines were treated every other passage with puromycin and assessed continuously for GFP expression by fluorescence microscopy.

#### Enzymatic Activity

Proteins immunoprecipitated by KDM6A antibody or by non-specific IgG (negative control) were measured for H3K27-directed KDM6 enzymatic activity using a chemiluminescent assay kit (50615, BPS Bioscience, Mira Mesa, San Diego, USA) following manufacturer’s instructions. Negative controls samples were incubated with 5ng/mL GSK-J4 dissolved in DMSO (S7070, Selleckchem, Houston, TX, USA) and compared to vehicle-treated samples.

### Quantification of Golgi apparatus morphology

Images were acquired using an Agilent BioTek Cytation C10 confocal microscope. Cells were randomly chosen for measurement (n=50-150 cells per group). ImageJ was used to quantify the Golgi apparatus (GM130) and nucleus (DAPI) area as well as distance from the nucleus to the furthest point of the Golgi apparatus. Normalized area was determined by dividing the area of the Golgi apparatus by the area of the nucleus for the same cell. Distance was measured by determining the perpendicular length from the edge of the nucleus to the furthest edge of the Golgi apparatus. Golgi elements were determined from maximum intensity projections z stacks of deconvoluted and Li thresholded particle analysis (Image J, NIH; https://image.nih.gov/ij/).

### Mass Spectrometry

Proteomics data for subcellular fractions deposited in MassIVE (MSV000094130) was reanalyzed [18]. This dataset contains liquid chromatography - ion mobility - tandem mass spectrometry (LC-IMS-MS/MS) data of fractionated MCF7 cells (cytoplasmic, membranous, soluble nuclear, chromatin-bound nuclear, and cytoskeletal). Four sample-cleanup techniques were used to remove contaminants following fractionation and protein extraction, including single-pot, solid-phase-enhanced sample preparation (SP3),[19] solvent precipitation, single-pot, solid-phase-enhanced sample preparation (SP4),[20] SP3 with a detergent-assisted digestion (SP3(DA)), and SP4(DA) [18]. Three sample preparation replicates and three technical replicates were acquired for each cleanup technique for each of the five subcellular fractions, totaling 180 total LC-MS/MS runs. Additional information on protein digestion with Trypsin/LysC and LC-IMS-MS/MS (Instruments: Waters NanoACQUITY UPLC connected to a Waters Synapt G2-S Mass Spectrometer) can be found in our prior reference [18].

### Searching for MLL Compass-like Compass Member Identifications

We created a custom database by searching for MLL Compass-like Complex members in Uniprotkb. We searched for ‘WDR5’, ‘RBBP5’, ‘DPY30’, ‘ASH2’, ‘UTX’, ‘KDM6A’, ‘KMT2D’, ‘PTIP’, ‘PAI’, and ‘NCOA6’ in Uniprotkb then filtered the results to include only proteins from humans. This custom database was created November 2023 and includes 383 proteins (deposited in MassIVE). For each subcellular fraction, data runs from all four sample-cleanup techniques were imported into Progenesis QI for Proteomics (QIP) database searching software (v 4.2). The database searching process included importing data into Progenesis QIP, correcting the mass-to-charge ratios for Glu-1-Fibrinopeptide B lockmass, and enabling an ion accounting workflow. Search parameters consisted of peptide and fragment tolerances: automatic, enzyme specificity: cleavage of the C-terminus of lysine and arginine (/K-\P/R-\P), maximum ion charge: 20, missed cleavages: maximum 1, run alignment: automatic, peak picking sensitivity: automatic, maximum protein mass: 250 kDa. Modifications included fixed carbamidomethylation at cysteine, variable oxidation at methionine, and variable phosphorylation at serine, threonine, and tyrosine. A false discovery rate of <1% was used by iadbs.exe in Progenesis QIP creating randomized sequences “on the fly”; and ion matching requirements were set to 5 fragments/protein, 2 fragments/peptide, and 1 peptide/protein. Relative quantitation using Hi-3 with protein grouping was used.

### Searching for KDM6A Isoform Identifications

Progenesis QIP outputs protein groups, which are proteins identified together based on shared peptide identifications. Grouping protein identifications makes it challenging to determine sequence coverage maps for specific isoforms. To address this, we used Protein Metrics PMi-Byonic v3.6.0 (Cupertino, CA) to identify KDM6A isoforms in our data. In our preliminary results, in searching our data in Progenesis QIP against the human reference proteome (UP000005640), SP4(DA) enabled the highest detection of KDM6A, based on abundance. Therefore, SP4(DA) runs were searched by PMi-Byonic to identify KDM6A isoforms.

We created another custom database. ‘KDM6A’ was input into the Uniprotkb search engine (https://www.uniprot.org/), and on the ‘Advanced’ tab, additional filtering included setting the organism to ‘human’ and the gene name to ‘KDM6A’ or ‘UTX’. This custom database was created in August 2023 and included 29 isoforms of KDM6A (deposited in MassIVE). The following search conditions were used: precursor mass tolerance: 10 ppm, digestion cleavage: C-terminus of lysine and arginine residues with a semi-specific digestion specificity, recalibration lock mass: Glu-1-fibrinopeptide B (785.84206 m/z), missed cleavages: maximum of 1, false discovery rate: 1% with Byonic adding decoys and common contaminants to the protein database, possible modifications: fixed carbamidomethyl at cysteine (common2), variable oxidation at methionine (common 2), and variable phosphorylation at serine, threonine, and tyrosine (rare2). The isoforms detected were analyzed by confidence scores and the number of peptides matched to a protein.

All search results have been deposited into the MassIVE data repository with accession number MSV000096748.

### Statistical Analysis

Statistical analysis was conducted on GraphPad Prism (v11.0.2) using indicated tests.

## RESULTS

### KDM6A is lost in the nucleus but retains co-localization with the Golgi apparatus upon EMT

We previously showed that KDM6A protein expression is suppressed upon five days of TGF-β-induced EMT and restored upon five days of withdrawal of TGF-β, associated with EMT reversion [15]. Nevertheless, we noted that expression of KDM6A was not completely lost. To determine how EMT affects protein expression levels of KDM6A and its non-nuclear protein levels, we recapitulated this work in MCF10A cells, a non-transformed epithelial breast cell line, by treating cells with TGF-β over the course of 10 days then removing treatment for five days (Fig. 1A). We assayed for expression of KDM6A and its paralog KDM6B. Coinciding with our prior observations, KDM6A decreases upon TGF-β treatment and recovers upon TGF-β withdrawal (Fig. 1A). Interestingly, KDM6B increases upon TGF-β treatment and retains an elevated expression level despite TGF-β withdrawal (Fig. 1A). To determine the impact of changes in KDM6A and KDM6B expression on H3K27me3-directed demethylation activity, we performed an enzyme activity assay on whole cell lysates from MCF10A cells treated, for different durations of time, with TGF-β. Contrary to our expectation, total H3K27me3-directed demethylation activity was unchanged during the time course of TGF-β treatment and withdrawal (Fig. 1B), indicating that KDM6A and KDM6B may compensate for each other, or that diminished protein levels are not necessarily indicative of changes in enzymatic activity.

**Figure 1.**
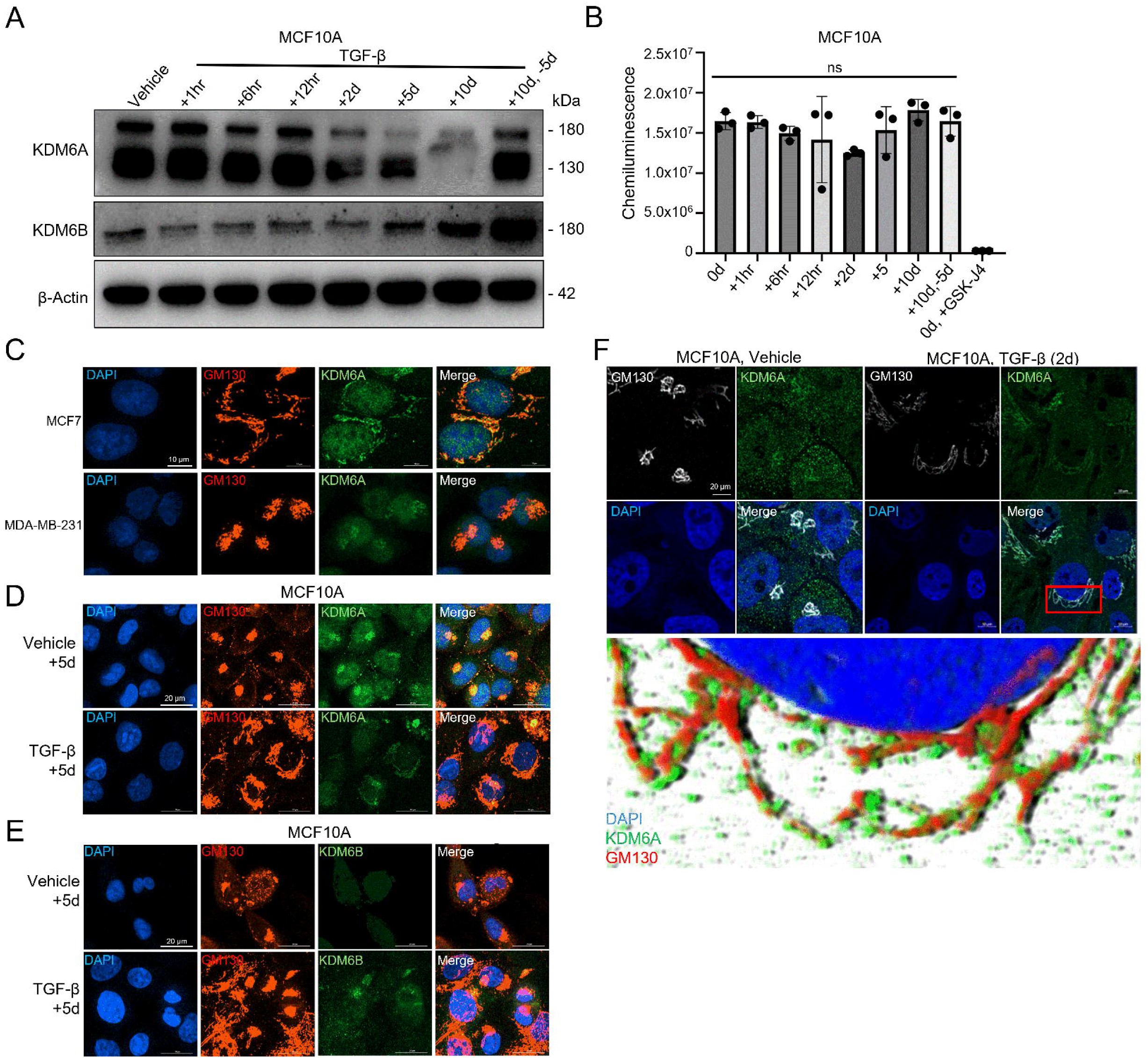
KDM6A and KDM6B localization to the Golgi apparatus. (A) Westen blot of MCF10A cells treated with 5.0 ng/mL TGF-β over the indicated time course and immunoblotted for the indicated proteins. β-actin was used as a loading control. (B) H3K27me3-directed activity of MCF10A whole cell lysates treated with TGF-β over the indicated time course was measured via activity assay. Addition of GSK-J4 served as a negative control. n = 3 cell culture replicates. Statistical significance tested using ordinary one-way ANOVA. (C) Confocal micrographs of MCF7 and MDA-MB-231 cells for KDM6A, GM130, and DAPI showing changes in Golgi apparatus morphology and KDM6A localization. Scale bar is matched for each image at 10 μm. (D) MCF10A cells treated with TGF-β or vehicle for five days immunostained for KDM6A, GM130, and DAPI. Scale bar is matched for each image at 20 μm. (E) MCF10A cells were immunostained for KDM6B, GM130, and DAPI after treatment with TGF-β for 5 days or vehicle. Scale bar is representative for each image: 20 μm. (F) MCF10A cells treated with TGF-β or vehicle for two days and immunostained for Golgi apparatus (GM130, white), KDM6A (Green), and counterstained for the nucleus with DAPI (blue). Scale bar: 20 μm. Zoomed-in image section is indicated by the highlighted box and was color enhanced. DAPI: blue, KDM6A: green, GM130: red.

We next observed the protein localization changes of KDM6A and KDM6B in this model of EMT, and in breast cancer cell lines with distinctive EMT phenotypes. Immunofluorescence was conducted on MCF7 and MDA-MB-231 cells, which express either E-cadherin or vimentin proteins (Sup. Fig. 1) and which are representative of the EMT(-) and EMT(+) cell states, respectively. Consistent with our prior results, MCF7 cells express high levels of KDM6A in the nucleus in contrast with MDA-MB-231 cells (Fig. 1C, Sup. Fig. 1). However, we note that, in addition to its nuclear localization, KDM6A is peri-nuclear in both cell lines and co-localizes with GM130, a marker of the Golgi apparatus (Fig. 1C). Further, we note that Golgi apparatus morphology changes with cell state, with Golgi in MCF7 cells adopting a dispersed pattern and Golgi in MDA-MB-231 cells forming a compact morphology.

As MCF7 and MDA-MB-231 cells differ in many respects besides EMT state, we assayed for KDM6A and KDM6B localization and Golgi morphology in TGF-β treated MCF10A cells as a model of inducible EMT. KDM6A but not KDM6B is lost in the nucleus upon TGF-β treatment (Fig. 1D-E). KDM6B expression increased upon TGF-β treatment and a portion of this protein was observed to co-localize with GM130. To more closely examine the co-localization of KDM6A and GM130, we analyzed images captured from cells treated with TGF-β for 40 hours using Airyscanning for super-resolution. As expected, KDM6A was detected co-localizing with GM130 in both vehicle and TGF-β treated cells (Fig. 1F). KDM6A enrichment at the Golgi apparatus ribbons is clearly visible (Fig. 1F-inset). Notably, TGF-β induces an apparent change in Golgi apparatus morphology from compacted to dispersed (Fig. 1D-F). To ensure that these observations were not influenced by choice of primary or secondary antibody, we conducted further tests in which fluorophores of the antibodies were varied. Altering the fluorophore did not impact the apparent Golgi localization of KDM6A; nor did a non-specific IgG primary antibody mimic this result (Sup. Fig. 2). Thus, our data indicate that Golgi apparatus morphology is influenced by epithelial-mesenchymal cell state and that KDM6A and KDM6B protein levels are localized to the Golgi apparatus.

### KDM6A isoforms are differentially expressed dependent on cellular compartment

To substantiate our observations made using immunofluorescence, we next turned to biochemical subcellular fractionation as a means to separately analyze nuclear and non-nuclear KDM6A. MCF7, MDA-MB-231, and MCF10A cells were subjected to subcellular fractionation to isolate cytoplasmic (C), membranous (M), soluble nuclear (SN), and chromatin-bound (CB) protein fractions for analysis via western blot.

As expected, KDM6A, KDM6B, and other protein members of the CLC (WDR5 and ASH2L) were consistently observed in the SN fraction in all three cell lines (Fig. 2A-C). Interestingly, KDM6A was detected in the M fraction of MCF7 and MDA-MB-231 cells while KDM6B was detected in the M fraction of MDA-MB-231 cells. For MCF10A cells, KDM6A and KDM6B were detected not only in the SN fractions, but also in the M and CB fractions (Fig. 2A-C). Importantly, nuclear transcription factor SP1 was detected in the SN fraction but not in the M fraction indicating a separation between these two sub-cellular compartments. To our surprise, both KDM6A and KDM6B proteins were detected at multiple molecular weights and the isoforms differed in their localization to sub-cellular compartments (Fig. 2A-C). Full-length KDM6A is detected near 180 kDa and could be observed in all sub-cellular fractions of MCF7 and MCF10A cells and the SN fraction of MDA-MB-231 cells (Fig. 2A-C). At least two smaller isoforms at the apparent molecular weights of about 80 and 115 kDa are observed in the M and SN fractions. In contrast, full-length KDM6B, is observed in the M fraction of MDA-MB-231 cells while smaller isoforms were detected in the SN fraction of both cell lines (Fig. 2A/B). In MCF10A cells, full length KDM6B was detected in the SN and CB fractions while the smaller isoform was detected in the C and M fractions (Fig. 2C). Consistent with established diverse cellular roles, WDR5 and ASH2L were both detected in the M fraction of MCF7, MDA-MB-231 and MCF10A cells.

**Figure 2.**
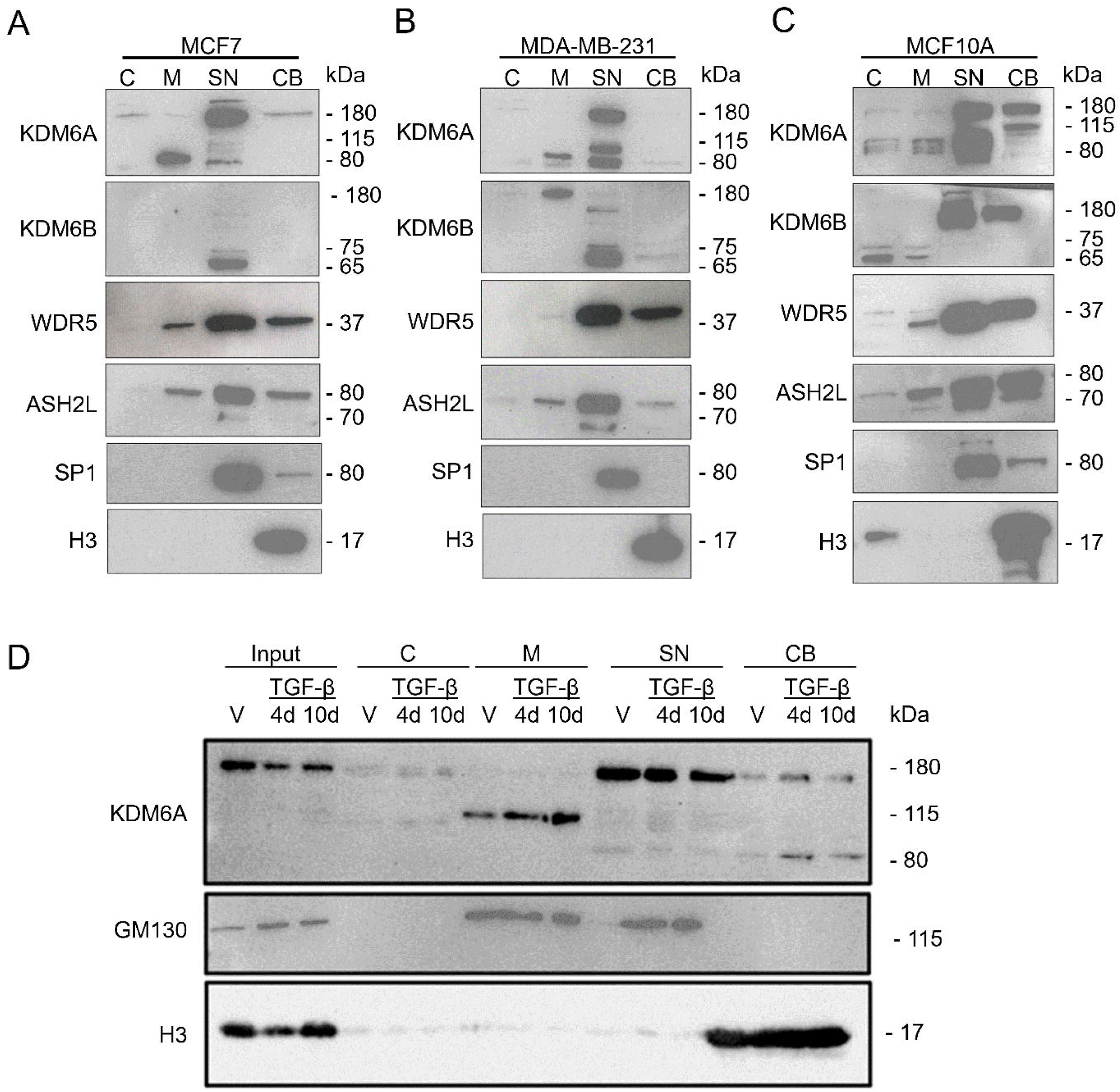
KDM6A isoform abundance differs dependent on subcellular localization and cell line. Western blot of MCF7 (A), MDA-MB-231 (B), and MCF10A (C) cells fractioned into subcellular fractions (C: Cytoplasmic M: Membranous SN: Soluble Nuclear CB: Chromatin Bound) and probed for the indicated proteins. (D) Western blot of fractioned TGF-β treated MCF10A cells for indicated proteins.

To determine if epithelial-mesenchymal state determines changes in KDM6A of different sub-cellular compartments, we treated MCF10A cells with TGF-β in a time course and conducted fractionation (Fig. 2D). In agreement with prior observations, KDM6A protein decreases upon EMT induction in the input and the SN fractions. KDM6A of the M fraction matches the smaller isoform, but more interestingly, membranous KDM6A appears to increase in relative abundance over the course of TGF-β treatment. Together these indicate that membranous KDM6A is of a shorter isoform than the full length KDM6A and is differentially expressed during EMT compared to the full length KDM6A localized in the canonical soluble nuclear location.

### Membranous-localized KDM6A is enzymatically active and contains the Jumonji C domain

As demonstrated above, KDM6A is expressed as proteins of various molecular weights. To better understand the non-full length isoforms of KDM6A, we analyzed data on known splice isoforms adopting the nomenclature of Fotouhi et al. (Fig. 3A). [21] While 8 of the 9 splice isoforms contain the catalytic (JmjC) domain one isoform, encoding a protein with a predicted molecular weight of 68.6 kDa, does not (Fig. 3A). While the short isoform from the M fractions ran at an apparent molecular weight between 80 and 115 kDa, it is nevertheless possible that this protein isoform is the shortest form of KDM6A, lacking a JmjC domain. Therefore, in order to determine if the membranous fraction possesses enzymatically active KDM6A, we performed an enzymatic activity assay following immunoprecipitation (IP) of KDM6A from sub-cellular fractions (Fig. 3B-C). IP of KDM6A from both the M and SN fractions yielded protein with similar enzymatic activity against a synthetic H3K27me3 peptide (Fig. 3B). This activity was sensitive to GSK-J4, a KDM6 family inhibitor and was not detected in IgG control samples (Fig. 3B). Interestingly, the isoform of KDM6A recovered by IP from the M fraction was the full-length isoform which was detected by western blot in the M fraction in MCF7 and MCF10A cells, though at a lesser intensity than the shorter isoform (Fig 2A).

**Figure 3.**
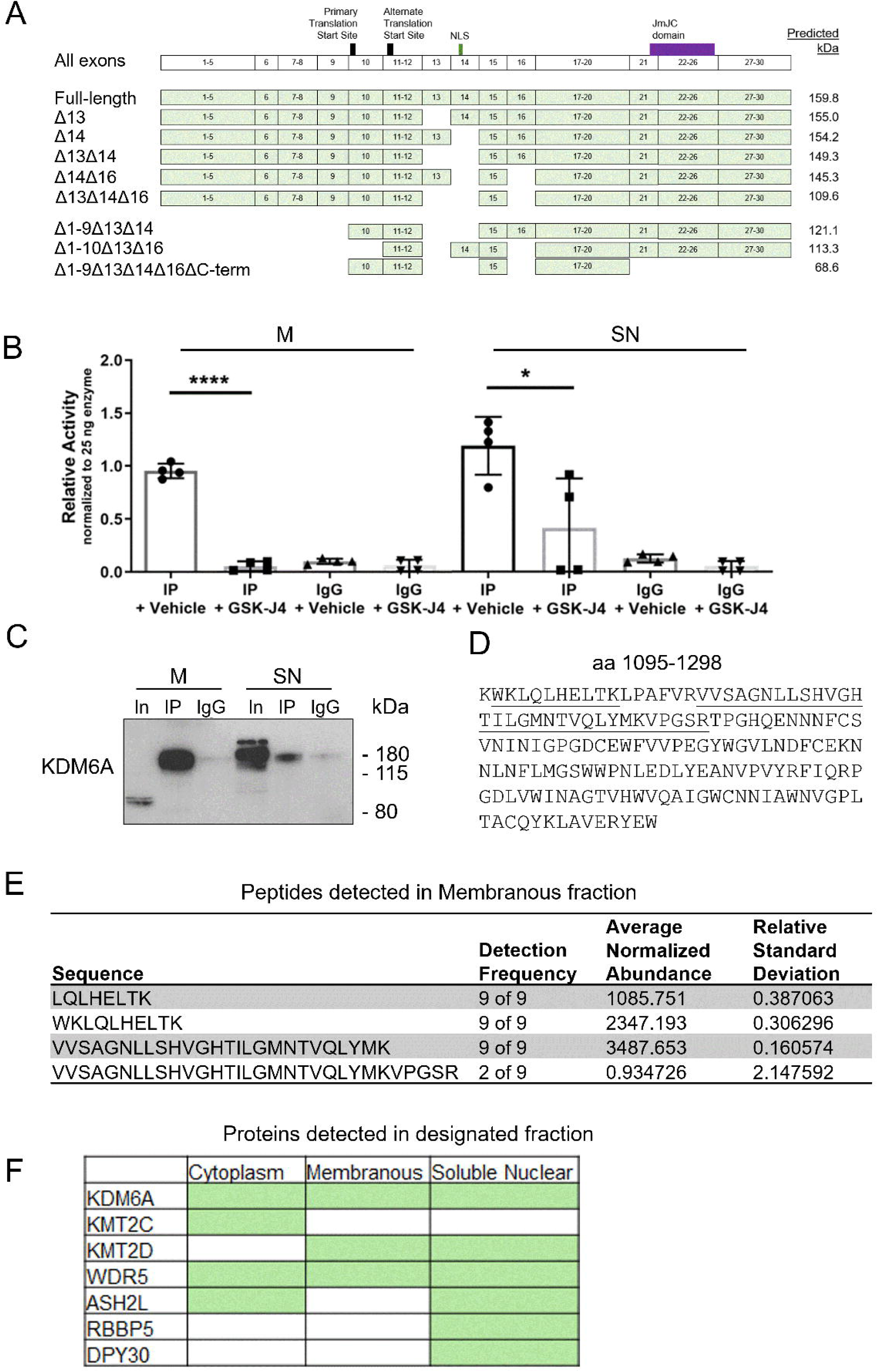
Non-nuclear KDM6A is enzymatically active outside its canonical location in MCF7 cells. (A) KDM6A isoforms with exon members and predicted protein size. Top-most row indicates the canonical and alternative translation start sites, the nuclear localization sequence (NLS), and the location of the Jumonji-C domain. (B) KDM6A activity assay conducted on immunoprecipitated KDM6A from the membranous and soluble nuclear fractions. n = 4 immunoprecipitant replicates Statistical significance tested unpaired t-tests between corresponding vehicle and GSK-J4 treated samples. (C) Western blot of KDM6A immunoprecipitated from MCF7 cell membranous and soluble nuclear subcellular fractions. (D) Amino acid sequence encoding the catalytic JmjC domain. (E) Peptides mapping to KDM6A detected in the membranous fraction via LC-MS/MS. (F) Detected peptides of indicated CLC members from mass spectrometric analysis of MCF7 subcellular fractions.

As a third method to characterize KDM6A outside the nucleus, we re-analyzed our previously published mass spectrometry from subcellular fractions of MCF7 cells data [18]. We were able to identify peptides mapping to known isoforms of KDM6A. We note that the membranous fraction contained peptides which map to the JmjC domain (Fig. 3D/E). Additionally, we tested if other members of the COMPASS-like protein complex could be detected in other protein subcellular fractions and indeed we were able to identify peptides that map to some, but not all the CLC members in the M and C fractions (Fig. 3F). In sum, while we cannot strictly conclude that the shorter isoform possesses enzymatic activity, available evidence suggests enzymatically active KDM6A is present in the M fraction, highlighting a potential role for KDM6A as a lysine demethylase outside the nucleus.

### Localization of other CLC members

As shown above, some CLC member proteins are detectable in the M fraction of MCF7, MDA-MB-231, and MCF10A cells (Figure 2A-C). Additionally, peptides mapping to KMT2D and WDR5, but not ASH2L, RBBP5 or DPY30, were detected in the M fraction of MCF7 cells (Fig. 3F) while western blots confirmed the presence of WDR5 and ASH2L (Fig. 2A). Biochemical separation of the M fraction is likely to include several organelles in addition to the Golgi apparatus. To further determine if CLC proteins co-localize with KDM6A at the Golgi apparatus, we performed co-immunofluorescent staining (Fig. 4).

**Figure 4.**
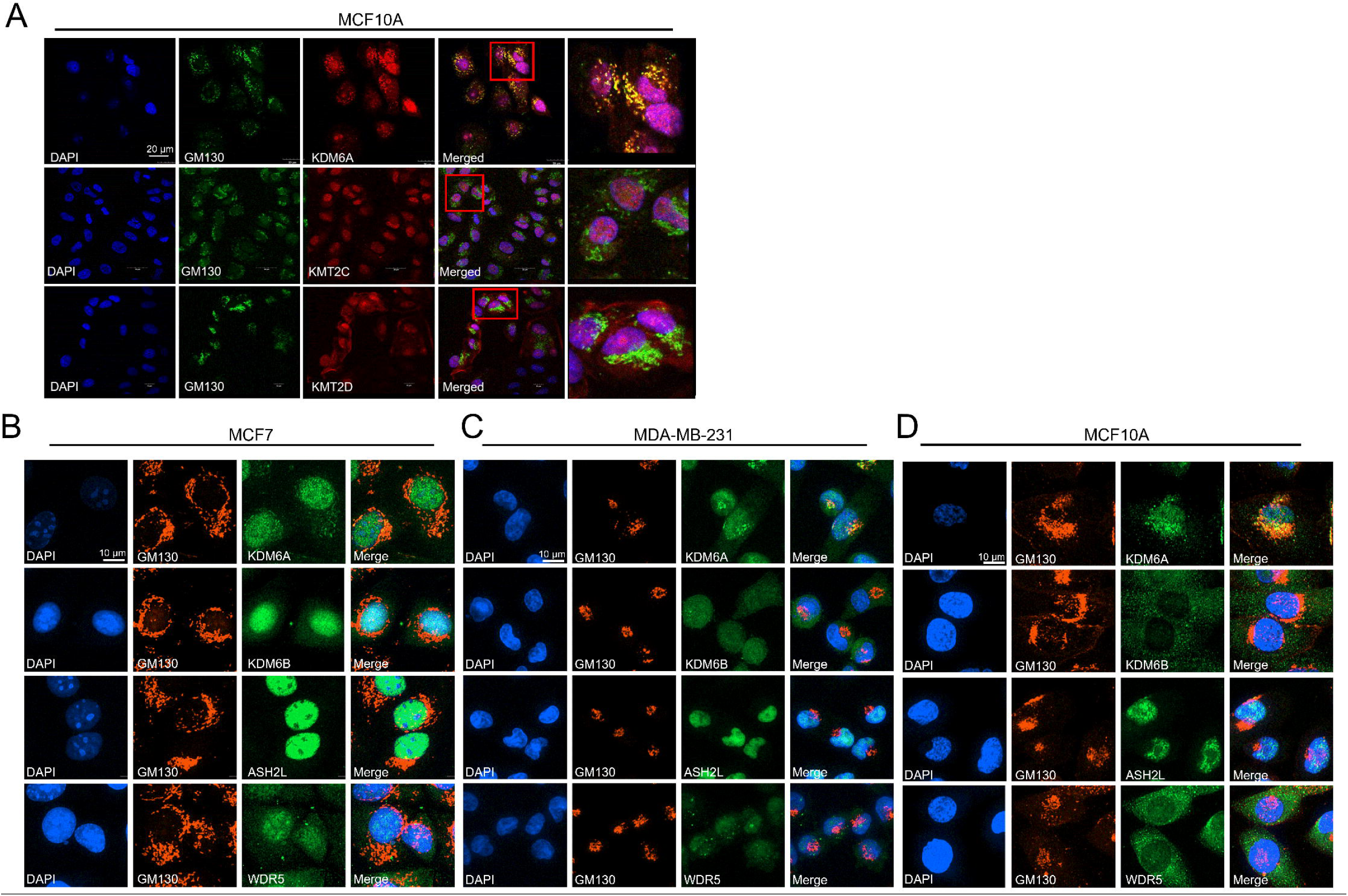
KDM6A is also found outside the nucleus co-localized with the Golgi apparatus. (A) Confocal micrographs of MCF10A cells stained for the following: KDM6A, GM130, and DAPI. Scale bar is matched for each image at 20 μm. MCF7 (B), MDA-MB-231 (C), and MCF10A (D) micrographs for the indicated COMPASS-like complex protein, GM130, and DAPI. Scale bar is matched for each image at 10 μm.

In MCF7, MDA-MB-231, and MCF10A cells we confirmed the colocalization of KDM6A with GM130 consistent with our prior observations (Fig. 4A-D). Co-localization of either KMT2C or KMT2D with GM130 was not observed for MCF10A cells, as both were primarily nuclear (Fig. 4A). Moreover, neither ASH2L nor WDR5 appeared to co-localize with GM130 at the Golgi apparatus in either MCF7 or MCF10A cells (Fig. 4B/C). As KDM6A does co-localize with the Golgi apparatus we next sought to determine if KDM6A expression or activity affects the morphology of the Golgi apparatus.

### KDM6A loss alters Golgi apparatus morphology similarly to EMT

We next sought to ascertain whether expression of KDM6A is relevant to Golgi apparatus morphology. Golgi apparatus morphology has been noted to be aberrant in many different cancer types such as lung, breast, and prostate cancers [22–25]. Indeed, data presented in Figure 1 indicate that two breast cancer cell lines exhibit distinct Golgi apparatus morphologies and that addition of TGF-β to MCF10A cells impacts morphology.

To determine if KDM6A expression is relevant to Golgi apparatus morphology, we knocked down KDM6A in MCF7 and MCF10A via lentivirus-drive short hairpin RNA (shRNA) (Sup. Fig. 3A). Cells were then stained for DAPI and probed for GM130 localization as a proxy for Golgi apparatus morphology. Images were analyzed for the change in relative Golgi apparatus area in addition to Golgi apparatus edge distance from the nucleus to gauge compaction (Fig. 5A). In MCF10A cells, loss of KDM6A expression caused a significant increase in both parameters (Fig. 5B/C). To determine if undergoing EMT similarly alters Golgi morphology, we next tested the effect of TGF-β on these parameters. As expected, consistent with our data in Figure 1, TGF-β caused a quantifiable change in relative Golgi apparatus area but did not significantly impact distance from the nucleus (Figure 5D/E).

**Figure 5.**
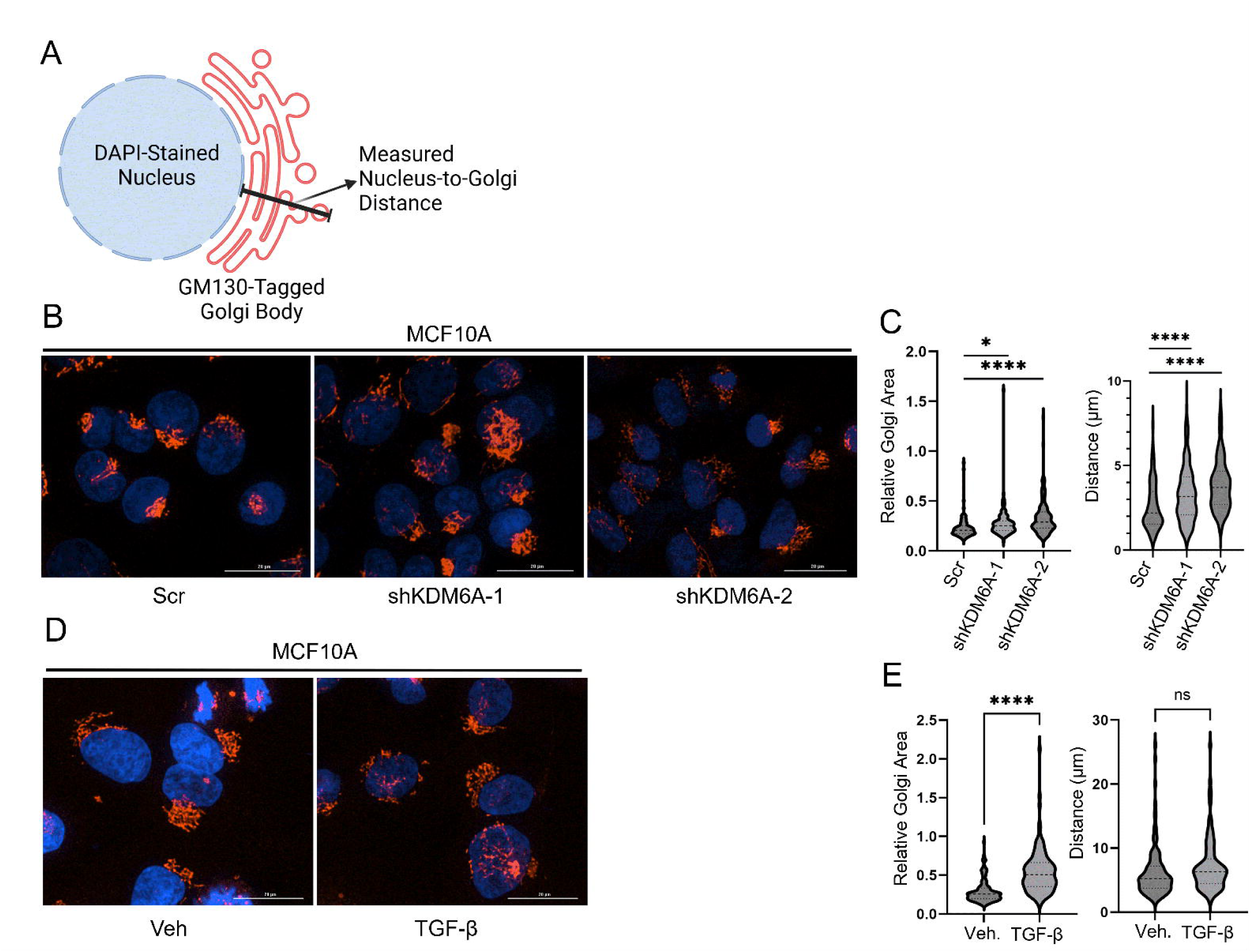
TGF-β phenocopies the effect of KDM6A loss on Golgi apparatus morphology in MCF10A cells. (A) Schematic illustrating measurement of Golgi distance, the distance from the nucleus perpendicular to the furthest Golgi body point. (B-E) Staining for nucleus (DAPI) and Golgi apparatus (GM130) was performed for (B) MCF10A cells in which KDM6A was knocked down by stable expression of either of two-targeted shRNAs (shKDM6A-1 & shKDM6A-2) and compared to a scrambled sequence control (Scr), and (D) cells which were treated with 5.0 ng/mL TGF-β or vehicle for 10 d. (C,E) Relative Golgi area (Golgi apparatus area / Nuclear body area) and nuclear-to-Golgi distance were measured per cell and analyzed. n > 90 cells per condition. *p < 0.05, **** p < 0.0001, ns = not significant.

We next tested the effects of these treatments in MCF7 cells. To our surprise however, KDM6A knockdown led to a decrease in relative Golgi apparatus area and decrease in the distance from nucleus to Golgi apparatus edge (Fig. 6A/B). These effects are diametrically opposed to our observations with MCF10A cells which exhibited enhanced distance from nucleus and relative Golgi area (Fig. 5B/C). Similarly, the impact of TGF-β on MCF7 cells was opposite of its impact on MCF10A cells (Fig. 6C/D). While the two cell lines responded differently to TGF-β and KDM6A knockdown, we note that the impact of TGF-β and KDM6A knockdown was the same within each cell line. Thus, KDM6A knockdown appears to phenocopy the effects of TGF-β on Golgi apparatus morphology.

**Figure 6.**
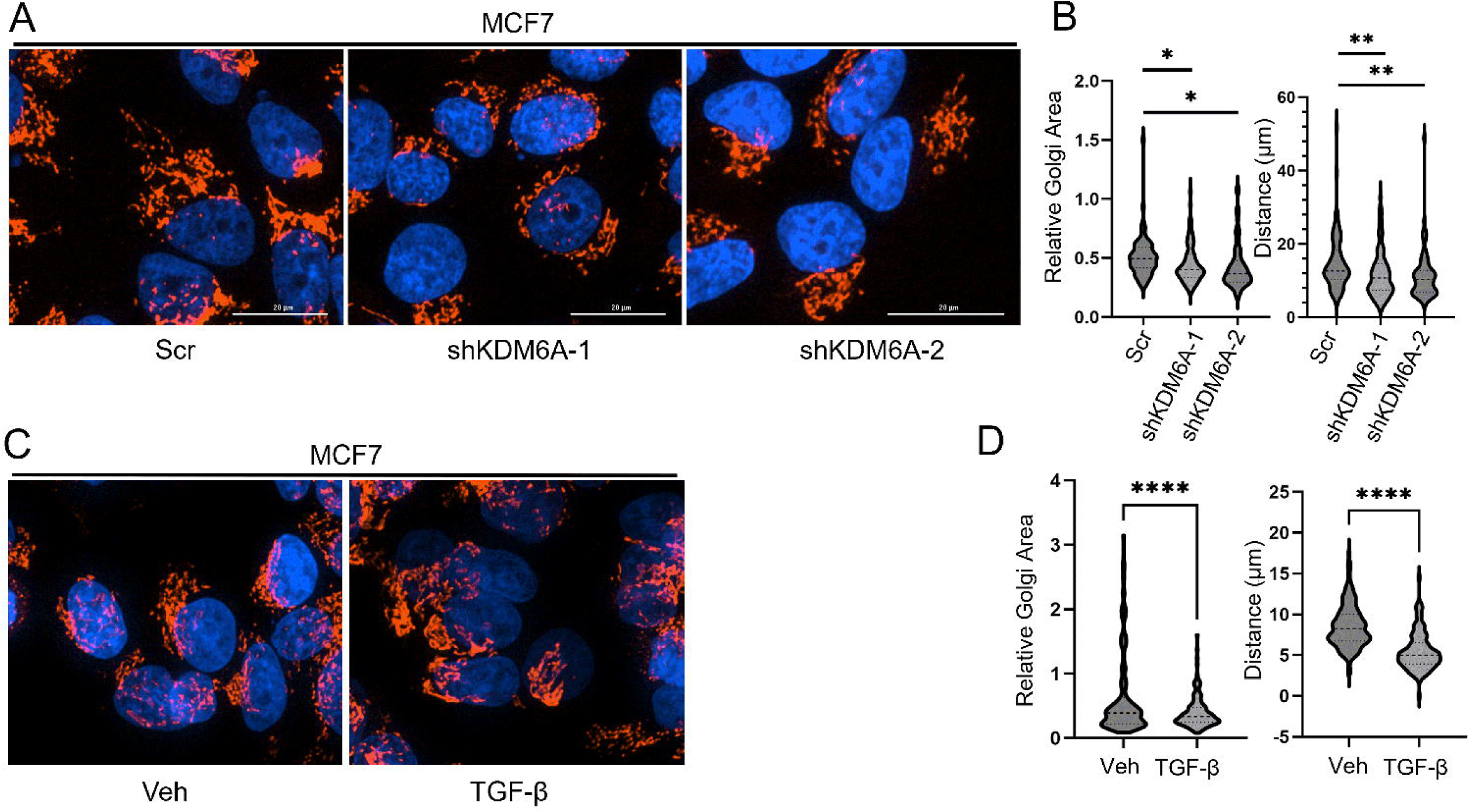
TGF-β phenocopies the effect of KDM6A loss on Golgi apparatus morphology in MCF7 cells. (A-D) Staining for nucleus (DAPI) and Golgi apparatus (GM130) was performed for (A) MCF7 cells in which KDM6A was knocked down by stable expression of either of two-targeted shRNAs (shKDM6A-1 & shKDM6A-2) and compared to a scrambled sequence control (Scr), and (C) cells which were treated with 5.0 ng/mL TGF-β or vehicle for 10 d. (B,D) Relative Golgi area (Golgi apparatus area / Nuclear body area) and nuclear-to-Golgi distance were measured per cell and analyzed. n > 69 cells per condition. *p < 0.05, ** p < 0.01, **** p < 0.0001.

We next considered that perhaps the distinctive responses of the MCF10A and MCF7 cells to either KDM6A knockdown or TGF-β treatment may be due to their non-transformed and transformed natures, respectively. To test this hypothesis, we also measured TGF-β induced Golgi apparatus changes in non-transformed and transformed human mammary epithelial (HMLE) cell lines: HMLE and HMLE cells transformed with the Ras oncogene (HMLER). In addition, we used HMLE-Twist cells as a positive control for EMT (Sup. Fig. 4). Unlike in the MCF10A, MCF7 cell paradigm, we found that in both HMLE and HMLER cells, TGF-β treatment significantly increased both relative Golgi area as well as Golgi distance. Thus, these two cell lines, non-transformed and transformed, responded most similarly to MCF10A cells. Interestingly, while there was not a significant difference in Golgi difference between HMLE and HMLE-Twist distance, the relative Golgi area was significantly decreased. As such, the Golgi apparatus morphology was altered in different directions dependent on EMT-induction pathway.

## DISCUSSION

KDM6A and KDM6B have both been shown to play roles in regulation of EMT to varying degrees dependent on tissue type. In particular, KDM6A has been shown to antagonize TGF-β-induced EMT in lung cancer cells while KDM6B promotes EMT in breast cancer cells [26] [27] [28] [29]. Our study reveals that KDM6A, a known epigenetic regulator which targets lysine 27 on histone 3, localizes to the Golgi apparatus despite nuclear loss during epithelial-to-mesenchymal transition. Through immunofluorescence we demonstrate that KDM6A is not only isolated to the nucleus, but also co-localizes with GM130, a marker or the Golgi apparatus while its paralog KDM6B does not (Figure 1). Furthermore, our data reveal that KDM6A, while lost in the nucleus upon EMT, is not lost in the Golgi apparatus, raising questions regarding its role outside the nucleus and if its potential impact on Golgi apparatus function. While KDM6A was previously shown to distribute within the cytoplasm, this is, to the best of our knowledge, the first characterization of Golgi localized KDM enzyme activity [30]. The identity of potential non-histone targets for KDM6A-mediated demethylation remains to be determined though such substrates have been identified for other JmjC-domain containing proteins including KDM4A [31].

KDM6A is expressed as a protein using multiple splice isoforms to generate distinct proteins, Fotouhi et al. found that the exon 12-17 region of KDM6A undergoes extensive splicing regulation. Notably, they observe that the two most abundant isoforms in MCF7 cells are the Δ14 and Δ14Δ16 isoforms which both lack exon 14 which includes the putative NLS [21]. Localization of KDM6A protein to the nucleus is thought to depend on its interaction with KMT2C or KMT2D within the CLC complex. As neither KMT2C nor KMT2D were localized to the Golgi apparatus, we hypothesize that KDM6A protein molecules not interacting with the CLC complex are then free to associate with the Golgi apparatus.

Golgi apparatus specific protein PAQR3 (progestin and adipoQ receptors member 3) has been found to control H3K4 methylation though reducing KMT2A methyltransferase activity through sequestering WRAD sub-complex members to the Golgi apparatus [32]. Further another Golgi specific protein PAQR11 has been found to mediate Golgi compaction initiated by epithelial-to-mesenchymal transition (EMT) [33] [16]. Although there is a link between the interaction of WRAD sub protein complex and Golgi specific proteins and their effect on KMT2A activity, the impact of this regulatory axis on KDM6A is undefined. While we did not observe co-localization of WRAD complex members to the Golgi (Figure 4), we cannot formally exclude the possibility that KDM6A localization there is linked to the CLC. Indeed, only WDR5 was consistently detected outside the nucleus, matching prior observations [34]. Other members of the CLC have also been found outside the nucleus such as KMT2C, which has been detected in the cytosolic fraction, although like KDM6A, its role outside the nucleus has not been established or defined [35].

The Golgi apparatus plays a critical role in cancer progression as it can act as a hub to support high metabolic demand and can allow for proliferation and metastatic capability of cancer cells. Golgi morphology is frequently altered in cancer cells, shifting from well-organized perinuclear ribbons in normal cells, to fragmented, dispersed or expanded, or compacted structures [36]. This change in Golgi morphology also promotes tumor progression, metastasis, and survival through contributions towards atypical protein trafficking and aberrant glycosylation [37] [25]. Golgi dispersal has been linked to DNA damage, tumorigenesis, and altered glycosylation while EMT is linked towards Golgi compaction through improved ribbon linking and cisternal stacking [38].

We observed that loss of KDM6A expression led to Golgi unpacking in the MCF10A cell line but Golgi compaction in the MCF7 cell line. These observations were consistent whether KDM6A was knocked down selectively or whether KDM6A protein was reduced via TGF-β treatment. We initially hypothesized that it may be the transformed/non-transformed aspect of these two cell lines which led to the different effects of KDM6A reduction. However, use of the isogenic HMLE vs HMLE-Ras cell lines did not reinforce this hypothesis. MCF10A are a basal epithelial cell line while MCF7 exhibit characteristics of the luminal epithelial phenotype [39] [40]. It may be that aspects of this distinct differentiation fate may impact the effect of KDM6A reduction on Golgi apparatus morphology. Furthermore, in HMLE cells, TGF-β causes Golgi fragmentation while Twist causes Golgi compaction. Thus, it is likely that the specific path of EMT influences the Golgi morphology change in addition to cell or cancer type and that the effects of TGF-β may be pleiotropic, causing additional impacts on the cell.

In conclusion, our study demonstrates that nuclear KDM6A, but not Golgi-localized KDM6A, is lost upon EMT induction. We reveal that Golgi-localized KDM6A is enzymatically active and contains the catalytic JmjC domain. Additionally, KDM6A loss affects Golgi apparatus morphology in a manner that is phenocopied by TGF-β induced EMT but which remains cell type specific. Furthermore, our findings indicate that KDM6A may have non-histone demethylation targets outside the nucleus.

## Acknowledgements and Funding

We thank Dr. Kelsey Johnson for initial experimental observations. We thank the Baylor University Molecular Biosciences Center for support during the course of this work. This work was supported by 1R15GM146188 to J.H.T. and by R35GM150464 to E.S.G.

## Author contributions

C.S.W., J.M.C., A.M.N., J.M., J.T.B., R.B., A.M.Z., K.P., H.L., and L.K.C. performed experimental work, conducted formal analysis, and curated data. D.D.S. and J.A.K., provided essential resources. E.S.G. and J.H.T. designed the study, supervised, and secured funding. C.S.W. and J.H.T. conceptualized the work and drafted the manuscript. All authors participated in editing the manuscript.

## Supporting Information

Figures S1-S4 provided as supporting information.

