## Supporting Data for "KDM6A is enzymatically active at the Golgi apparatus where it is necessary for maintenance of Golgi apparatus morphology"

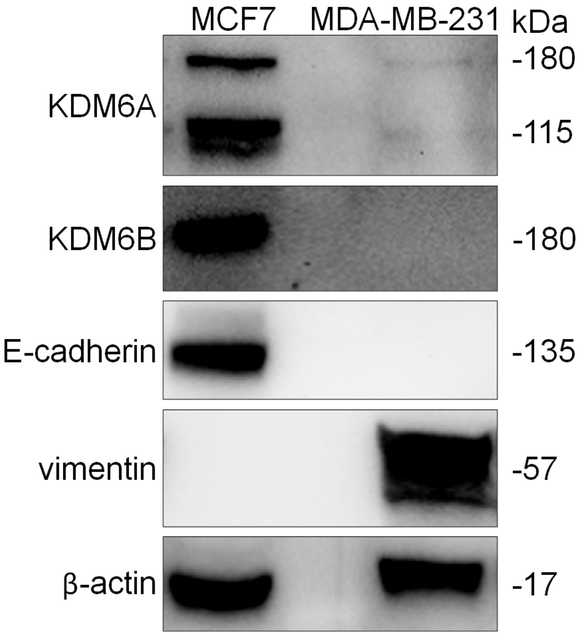


Supplemental Figure 1.

Expression of KDM6A and KDM6B as well as EMT markers in MCF7 and MDA-MB-231 cells. β-actin acts as a loading control. Center lane is not used

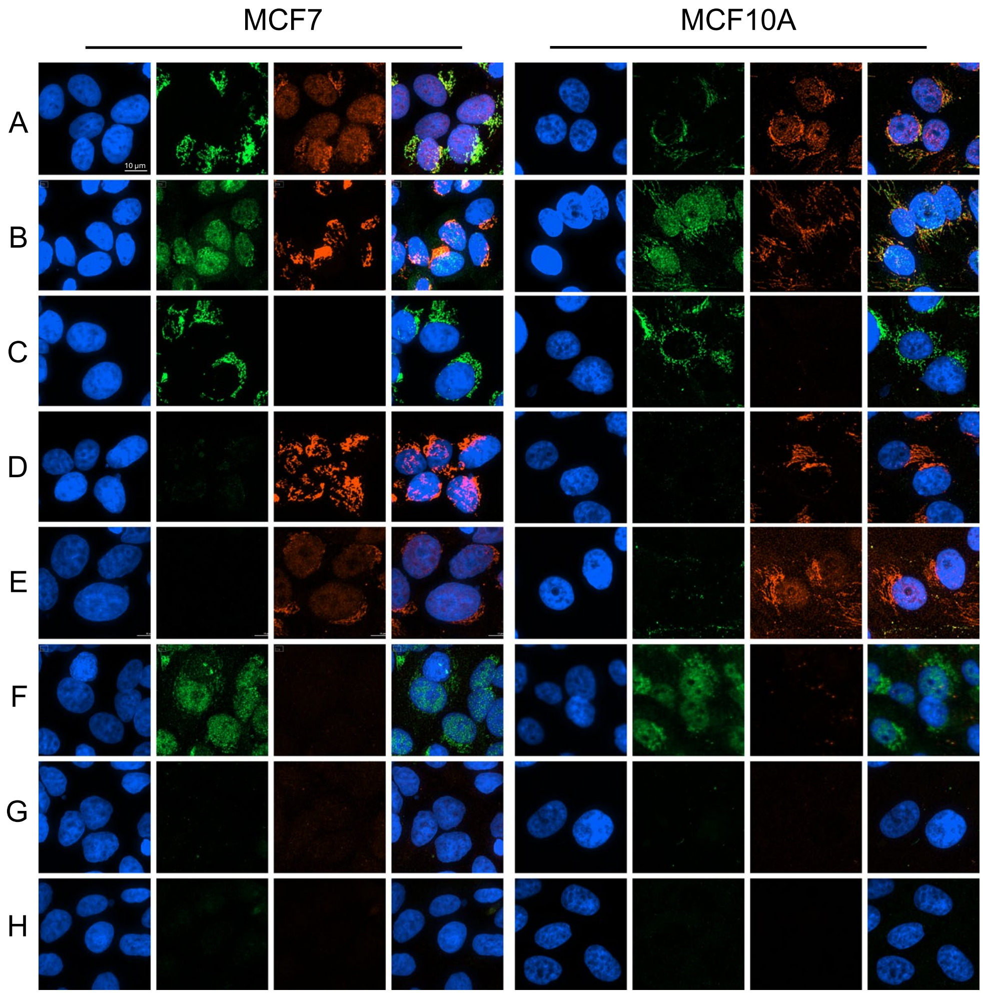


| | Green Channel | Red Channel |
|---|---|---|
| A | Mouse GM130 | Rabbit KDM6A |
| B | Rabbit KDM6A | Mouse GM130 |
| C | Mouse GM130 | Rabbit Normal IgG |
| D | Rabbit Normal IgG | Mouse GM130 |
| E | Mouse Normal IgG | Rabbit KDM6A |
| F | Rabbit KDM6A | Mouse Normal IgG |
| G | Mouse Normal IgG | Rabbit Normal IgG |
| H | Rabbit Normal IgG | Mouse Normal IgG |

Supplemental Figure 2. MCF7 and MCF10A cells stained with DAPI and immunostained for GM130 and KDM6A. Table represents the representative fluorescent color of the indicated protein. Scale bar is representative for each image: 10 μm.

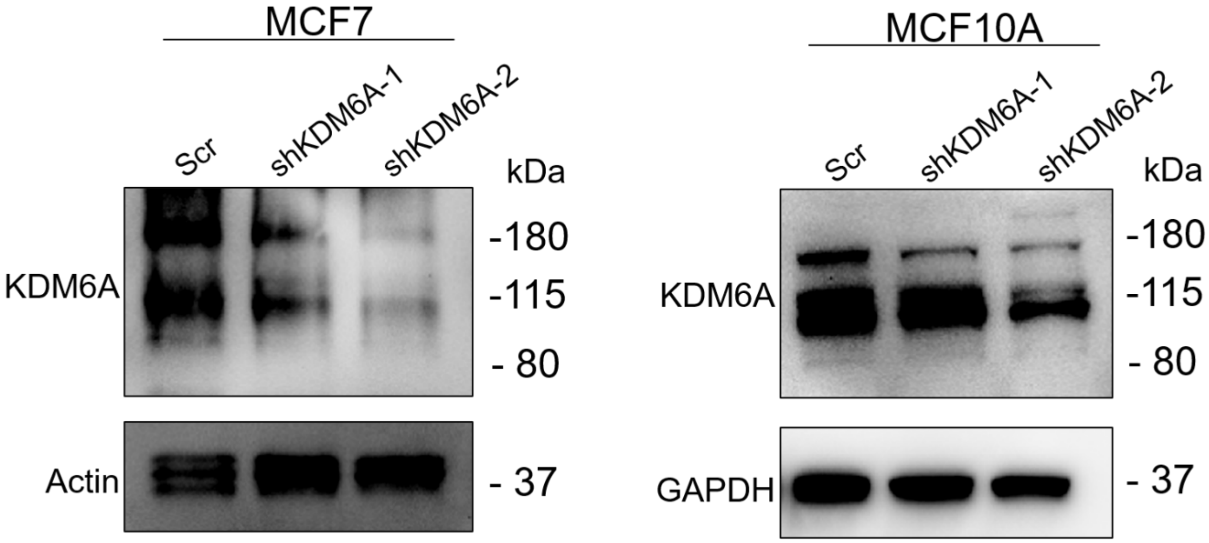


Supplemental Figure 3. Western blot of indicated proteins from indicated cell lines. Scr indicates non-targeted control shRNA. shKDM6A-1 and shKDM6A-2 are independent shRNA sequences both targeting KDM6A. GAPDH and β-actin were used as loading controls.

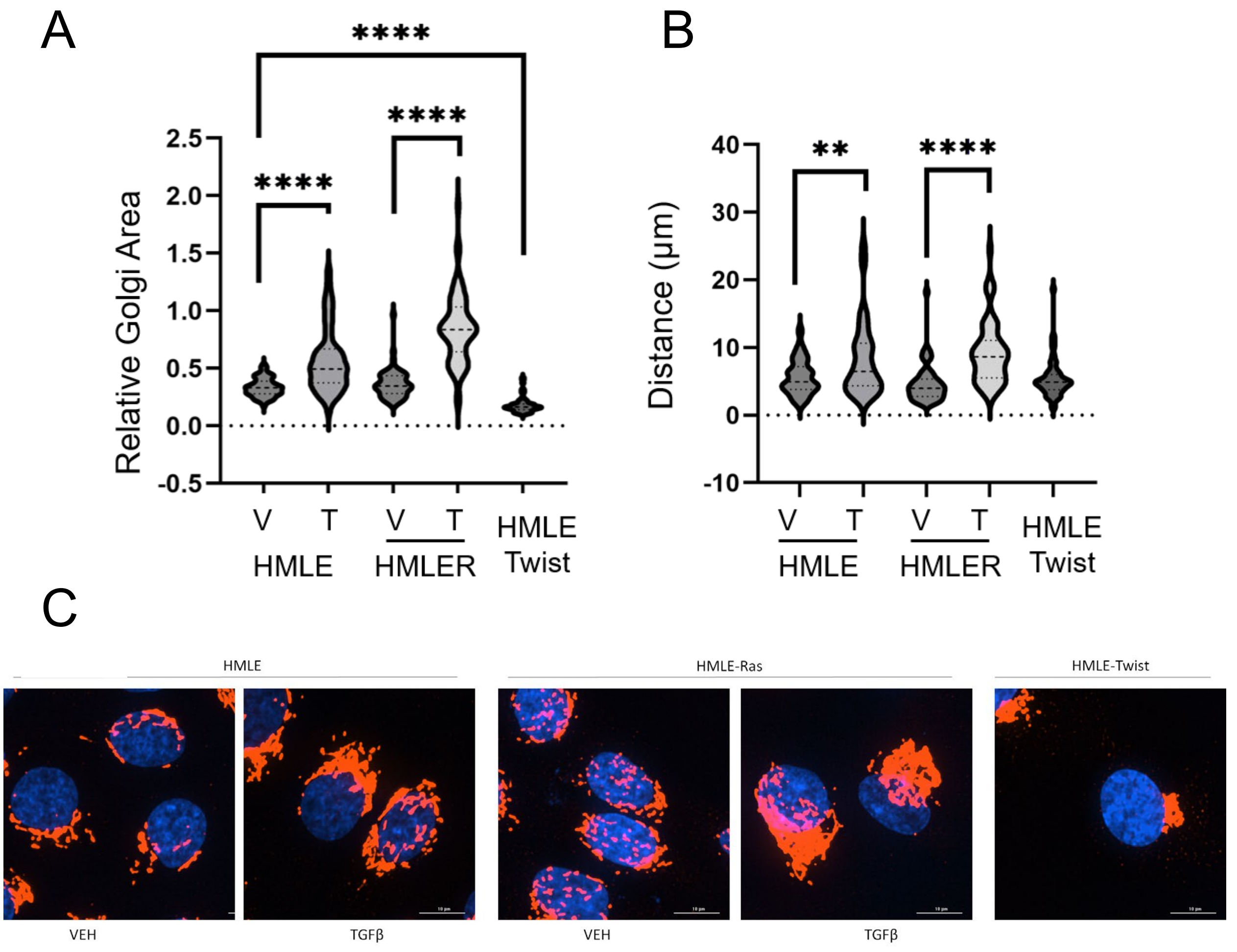


Supplemental Figure 4. Impact of TGF-β and Twist expression on Golgi apparatus morphology in mammary epithelial cells. (A,B). Relative Golgi area (Golgi apparatus area / Nuclear body area) and nuclear-to-Golgi distance were measured per cell and analyzed. n > 80 cells per condition. (C) Staining for nucleus (DAPI) and Golgi apparatus (GM130) was performed for indicated cell lines. Cell treatments consisted of 5.0 ng/mL TGF-β or vehicle for 48 h. **p<0.01; ****p<0.0001
